# Enhancing pre-defined workflows with *ad hoc* analytics using Galaxy, Docker and Jupyter

**DOI:** 10.1101/075457

**Authors:** Björn A. Grüning, Eric Rasche, Boris Rebolledo-Jaramillo, Carl Eberhard, Torsten Houwaart, John Chilton, Nate Coraor, Rolf Backofen, James Taylor, Anton Nekrutenko

## Abstract

What does it take to convert a heap of sequencing data into a publishable result? First, common tools are employed to reduce primary data (sequencing reads) to a form suitable for further analyses (i.e., list of variable sites). The subsequent exploratory stage is much more *ad hoc* and requires development of custom scripts making it problematic for biomedical researchers. Here we describe a hybrid platform combining common analysis pathways with exploratory environments. It aims at fully encompassing and simplifying the “raw data-to-publication” pathway and making it reproducible.

## 1 Introduction

Trees, rivers, and the analysis of next generation sequencing (NGS) data are examples of branching systems so ubiquitous in nature [1]. Indeed, numerous types of NGS applications (i.e., variation detection, ChIP-seq, RNA-seq) share the same initial processing steps (quality control, read manipulation and filtering, mapping, post-mapping thresholding etc.) making up the trunk and main branches of this imaginary tree. Each of these main branches subsequently gives off smaller offshoots (variant calling, RNA-, ChIP-and other “seqs”), that, in turn, split further as analyses become focused towards the specific goals of an experiment. As we traverse the tree, the set of established analysis tools becomes increasingly sparse and it is up to an individual researcher to come up with statistical and visualization approaches necessary to reach the leaves (or fruits) representing conclusive, publishable results. Consider transcriptome analysis as an example. Initial steps of RNA-seq analysis (in our tree allegory these are trunk and main branches), such as quality control, read mapping, transcript assembly and quantification, are reasonably well established. Yet completion of these steps does not produce a publishable result. Instead, there is still the need for additional analyses (progressively smaller branches of our tree) ranging from simple format conversion to statistical tests and visualizations. Thus every NGS analysis can in principle be divided into two stages. (1) The first stage involves processing of raw data using a finite set of common generic tools. This stage can be scripted and automated and also lends itself to building graphical user interfaces (GUIs). (2) The second stage involves a much greater variety of tools that need to be customized for every given experiment (in many cases there are no tools at all and custom scripts need to be developed). As a result it is not readily coerced into a handful of automated routines or generic GUIs.

The main motivation for this work was the development of a system where biomedical researchers can perform both stages of data analysis: initial steps using established tools and exploratory and data interpretation steps with *ad hoc* approaches. Merging both steps into a unifying platform will lower entry barriers for individuals interested in data analysis, significantly improve reproducibility of published results, ease collaborations, and enable straightforward dissemination of best analysis practices.

## 2 Results and Discussion

To avoid “reinventing the bicycle” in designing our platform, we first evaluated already existing systems that can be leveraged to fulfill our goals. For the first stage of the analysis we needed a system that exposes existing common tools through a unifying interface and makes computational infrastructure needed to perform large-scale analyses transparent to the user. There are several systems potentially satisfying these requirements including Genepattern [2], Mobyle [3], CyVerse [4], Galaxy [5], and GenomeSpace [6]. These systems allows users to utilize a large number of tools and workflows as well as record provenance ensuring reproducibility of analyses. However, these systems are only as useful as their set of featured tools, and do not aid in *ad hoc* data exploration. The number available choices for the second stage of the analysis (*ad hoc* exploration) is enormous. One can simply use scripting languages, relational databases, spreadsheet applications, or commercial packages to conduct data interpretation. One important feature we sought is the ability for a system to record analysis steps in order to make final outcome reproducible. In this regard two well established open environments designed specifically for reproducible data exploration stood out as *de facto* standards in scientific computing: IPython/Jupyter [7] and RStudio [8].

In the end we proceeded with Galaxy (due to its considerable user base http://bit.ly/gxyStats) as the underlying platform for management of data, tools, and infrastructure and Jupyter as an initial data exploration plug-in (at the time of writing RStudio has also been integrated and being tested). Galaxy is a web-based analysis environment that exposes tools using GUI, allows combining them into workflows, and is supported by software and hardware infrastructure suitable for analysis of very large multi-sample datasets. The benefit of Galaxy is that anyone with a web-browser can perform analyses in a straightforward manner without being concerned with how or where underlying software is executed. Jupyter (formerly known as IPython) is an interactive programming environment allowing reproducible data analysis with over 40 programming languages (such as Python, Julia, R, and others). It is built around the concept of Jupyter Notebook – a web application allowing to combine executable programming language code with visualization, and explanatory annotations in a single “live” document. The advantage of Jupyter is that there is essentially no limit on what one can do as supported languages and underlying libraries enable the full spectrum of data analyses. However, to be useful Jupyter requires programming and data management skills as well as access to computational infrastructure. Table 1 contrasts the pros and cons of the two platforms (Galaxy and Jupyter) and shows that their combination provides an almost “perfect” analysis solution for biomedical domain.

**Table 1.** Congruence between Galaxy and Jupyter as a function of their pros and cons.

| Feature | Galaxy | Jupyter | Galaxy/Jupyter |
| --- | --- | --- | --- |
| Low barrier of entry for a naive user | * |  | * |
| Versatility of available tools |  | * | * |
| Provenance tracking | * | * | * |
| Hardware backend for processing of large datasets | * |  | * |
| Attractive to experimentalists | * |  | * |
| Attractive to bioinformaticians and data scientists |  | * | * |

How can these two very dissimilar applications, Galaxy and Jupyter, work together in practice? In Galaxy, datasets corresponding to each step of analysis are recorded in the history as “history items” (right panes of the interface in Fig. 1A and 1B). Once an analysis reaches the point where there are no tools available for the next step, it is time to switch to Jupyter. This is done by clicking a button adjacent to the dataset, which will start an isolated instance of Jupyter (or other so called Interactive Environments, such as RStudio) directly within Galaxy interface. This instance interacts with Galaxy’s Application Programming Interface (API) using custom methods for transferring data back and forth from Galaxy’s history. Jupyter’s “notebooks” allow *ad hoc* analyses to be recorded automatically, providing the utmost level of reproducibility during data exploration. After performing analyses the user can save the notebook as a Galaxy history item that can be downloaded and used as a template for a new tool, it can be re-run with changed parameters on different datasets and like any other history item it can be shared with other Galaxy users. Additionally, in the future it will be possible to export the code, aiding in rapid development of galaxy tools. It can also be converted into a PDF documenting details of the analyses. Here we used this functionality to generate Supplementary Files 1, 2, and 3 corresponding to the three examples described below.

**Figure 1.**
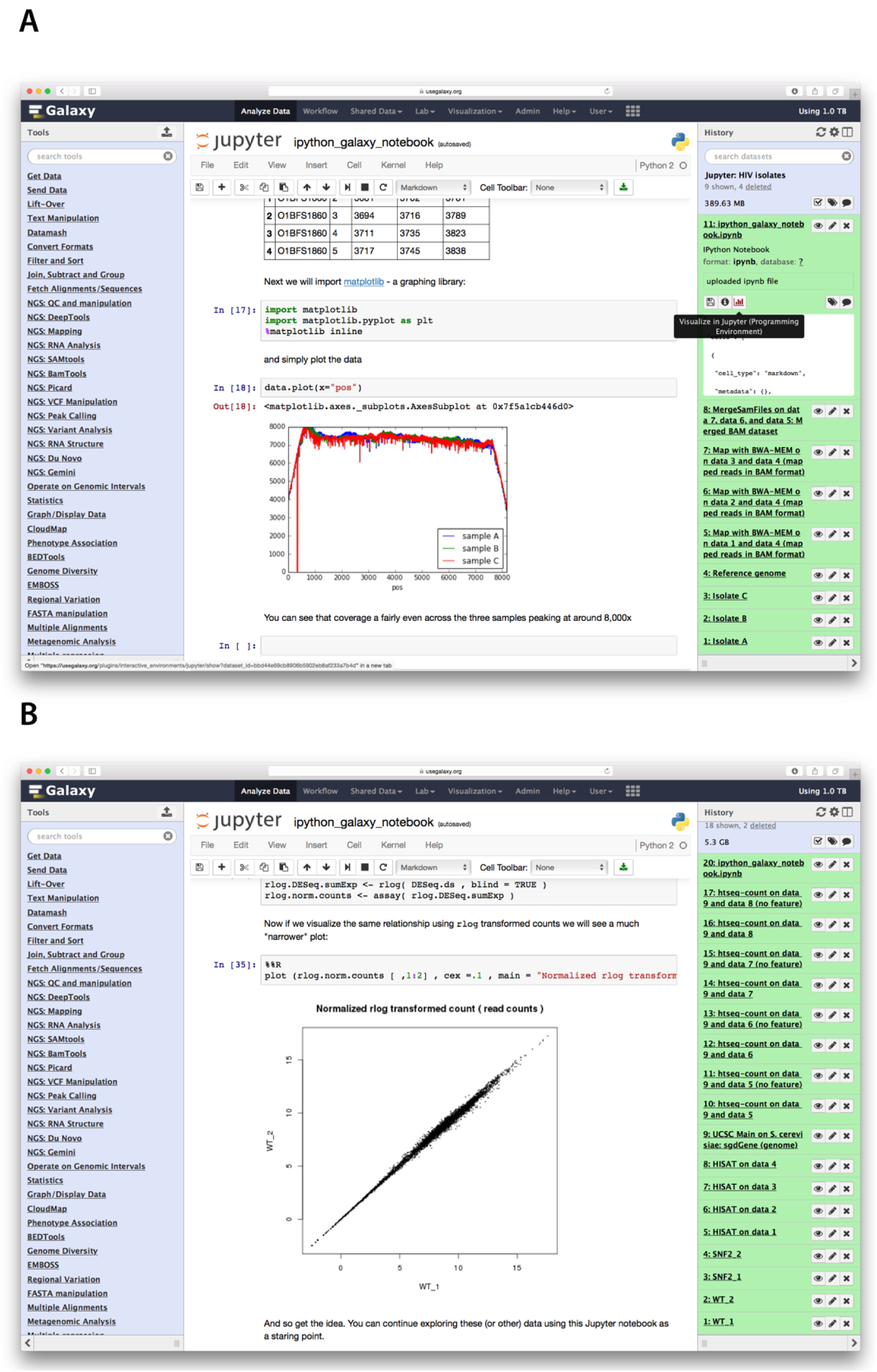
Overview of steps involved in performing analyses outlined in Examples 1 and 2. **A.** Example 1. Right (green) side of Galaxy interface is the history pane. The analysis begins with uploading three Illumina datasets (datasets 1-3) and a reference genome sequence (dataset 4). Datasets are mapped to the reference genome with bwa-mem (datasets 5-7) and read groups are assigned (datasets 8-10). This allows resulting BAM datasets to be merged into a single BAM file (dataset 11). At this point the Jupyter IE is launched. Lower part of the notebook is visible in the center pane showing the read coverage distribution for the three isolates (three different colors). **B.** A similar screenshot for Example 2. Here Illumina reads for two RNA-seq replicates from wild-type and snf2 knock-out are mapped against the Drosophila melanogaster genome (dm3) using HiSat split mapper. Next, HTSeq-count takes BAM datasets generated by HiSat and, using gene annotation for dm3 genome downloaded from the UCSC Table Browser (history dataset 9), computes per-gene read counts. These counts are then imported to Jupyter (Center pane) to perform normalization and variance shrinkage calculations using Bioconductor’s DESeq2 package.

To demonstrate the utility of Galaxy/Jupyter integration we devised three examples. In all three cases we break down the analysis in parts 1 (Galaxy) and 2 (Jupyter). For each example, part 1 involves processing and mapping of the sequenced reads. In the first example, we use a simple combination of command line tools and Python scripting language to plot read coverage across the HIV genome. In the second example, we leverage Python and R to normalize read counts and shrink variance in an RNA-seq experiment. Finally, in the most complex example, we perform data processing and replicate main summary figures from our previous study [9]. This third example demonstrates the capabilities of Galaxy and Jupyter to process large, multi-sample datasets. In this example Galaxy’s power is leveraged for mapping and processing of hundreds of datasets and Jupyter is used for the final interpretation and replication of published figures. These examples can be viewed in the supplemental PDFs, or they can be interacted with in a live Galaxy/Jupyter instance by pointing your browser to the links indicated below.

### 2.1 Example 1: Building a genome coverage plot

HIV-1 was re-sequenced from blood of a single individual across three time points with the ultimate goal of tracking nucleotide substitutions of the viral genome through time (simulated reads were generated for this example). After assessing the quality of the reads and mapping against the HIV-1 genome with bwa [10] within Galaxy, we wanted to visualize read coverage across each sample to decide if further analyses are warranted. However, the main public Galaxy server did not have a dedicated tool for this purpose. Normally the analysis will stop at this point and only by downloading data and analyzing them offline can one produce the coverage distribution graph needed in this case. Integration of Jupyter to Galaxy changes this. Fig. 1A highlights each step of this analysis resulting in the coverage distribution graph. The entire analysis can be seen in the Galaxy history accessible at http://bit.ly/ie-hiv.

### 2.2 Example 2: Normalizing read counts for an RNA-seq experiment

In this example we use a subset of RNA-seq data from a dataset published by Schurch et al. [11] (SRA accession ERP004763) consisting of 48 replicates of two *Saccharomyces cerevisiae* populations: wildtype and snf2 knock-out mutants. For simplicity we selected only two replicates for each wild-type and snf2 knock-outs. Here we first use Galaxy’s existing RNA-seq tools to map reads against the yeast genome using HiSat [12] and to compute the number of reads per gene region using HTseq-count [13] (Fig. 1B; see Galaxy history at http://bit.ly/rnaseq-jupyter). Datasets are then imported into Jupyter’s environment (cells 4 - 9; see Supplemental Document 2) where we first merge datasets into a single table by joining them on gene names using Python’s Pandas library (cells 10 - 12). We then proceeded to normalize the counts with DESeq2 [14] (cells 13 - 30) and assessed the effects of normalization and variance shrinkage on the data (cells 31 - 35; also see center pane of Fig. 1B).

### 2.3 Example 3. Estimating mitochondrial bottleneck in humans

In the third example we replicate the key analyses reported in a study of human mitochondrial heteroplasmy transmission dynamics, previously published by our group [9]. The overall goal of this study was to detect heteroplasmies (variants within mitochondrial DNA) and to trace their frequency changes across mother-child transmission events using primary sequencing data generated by [9] (mitochondria is transmitted maternally and heteroplasmy frequencies may change dramatically and unpredictably during the transmission, due to a germ-line bottleneck [15]). The first part of the analysis is performed using Galaxy’s mapping and variant calling workflow outlined in Fig. 2A. The goals of this part is to generate a preliminary list of sequence variants. The input data consists of ≈118 Gb of data corresponding to 312 fastq datasets (SRA accession SRP047378) derived from 156 samples 156 samples (39 mothers and 39 children, with two tissues analyzed per individual, each tissue generating two fastq datasets for the forward and reverse read sets, together resulted in the 312 original datasets; Fig. 2B, dataset 313). Using Galaxy, we combine all 312 datasets into a single entity, a dataset collection, in order to avoid repetitive tasks (see Galaxy history at http://bit.ly/jupyter-mt and Fig. 2B). The workflow maps the reads, performs de-duplication and extensive filtering of resulting BAM datasets as well as identifies variable sites. The workflow reduces sequencing reads to a 160 Mb data matrix with over 2.6 million rows containing variants for all 156 samples. Despite the fact that we have reduced the primary sequence data to a set of variable sites, this dataset hardly resembles an interpretable result. At this point, exploratory analyses must begin. Unfortunately it is also the point where users are forced to leave Galaxy, confounding efforts for reproducibility of the analysis. With Galaxy/Jupyter integration, this deficiency can be avoided. The second part of the analysis begins with starting a Jupyter notebook from inside the Galaxy interface (see Supplemental Document 3). The goal of the analysis is to estimate the size of the germ-line bottleneck and to detect the effect of maternal age on the frequency of heteroplasmic variants observed in children (for details see [9]). In performing this analysis we first check for even coverage across all samples (cell 24 of the notebook), and then we threshold the sites to retain a set of 181 variants with the minor allele frequency (MAF) above 1% that do not display strand bias calculated using Guo at al. formula [16] (cells 27 - 31). Next, we calculate the statistical significance of these sites using a Poisson distribution as in Li and Stoneking [17], which is based on the variability of a given position across all samples (cells 32 - 35). These two calculations (strand bias and statistical significance) are performed by implementing formulae from two publications directly in Jupyter. This is a convincing example of utility for such an environment: it is impractical to write dedicated tools based on these formulae in Galaxy because they will not be sufficiently general, yet it makes perfect sense to do it in Jupyter on an as-needed basis. The notebook is stored as a history item in Galaxy making the analysis permanent and perfectly reproducible (Fig. 2B, dataset 1417). The analysis proceeds further by screening for contamination based on heteroplasmy frequency distribution (large number of heteroplasmies with a tight frequency distribution is indicative of contamination [18]; cell 37) and tabulating the distribution of variants across individuals and tissues (cell 57) and features of the mitochondrial genome (cell 60). At this point we arrive to one of the main observations of the study (cell 67): the heteroplasmy frequency in children does not diverge between tissues (Fig. 2C; pane A compared to mother, pane B) as they underwent fewer mitotic segregations and were exposed to potential environmental effects for a shorter period of time. In addition, there is a stronger correlation in allele frequencies between two tissues of mother or a child than between mother and child for the same tissue (Fig. 2C; panes C and D) -a result of the germ-line bottleneck. Finally, using formulas from Millar, Hendy, and co-workers [19] we estimate the size of the germline bottleneck to be at 40 segregating units (cells 70 - 75; Fig. 2D). The final observation shown in cell 86 of the notebook highlights the positive correlation between the age of the mother and the number of heteroplasmic sites, which has potential implications for the higher rate in mtDNA diseases in children born to older mothers.

**Figure 2.**
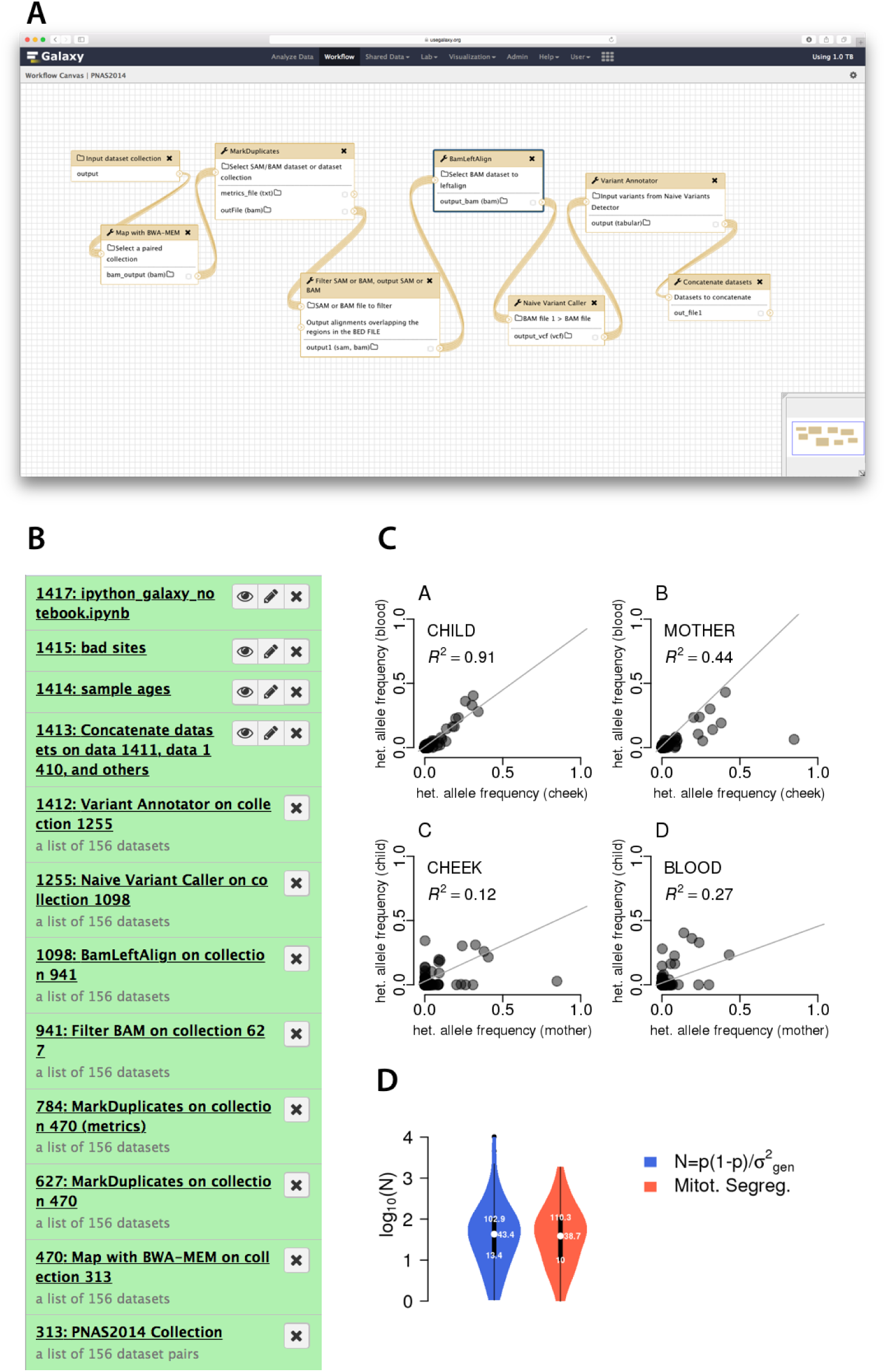
Re-analysis of data from [9] using Galaxy and Jupyter. **A.** Workflow used in the analysis. As an input the workflow takes a collection of paired illumina datasets and outputs an unfiltered list of variable sites. **B.** Galaxy history showing all steps of these analyses. It only contains 12 steps because we use dataset collections to combine multiple similar datasets into a small number of history entries. This significantly simplifies processing. For example, collection 313 contains all 312 paired-end Illumina datasets generated for this study. This allows us to deal with just one history item instead of 312. The next item in the history is a collection of BAM datasets generated by mapping each read-pair from collection 313 against human genome (hg38) with bwa-mem. These BAM datasets are de-duplicated (collections 627), filtered (by only retaining reads mapping to mitochondrial DNA, with mapping quality of 20 or higher, and mapped in a proper pair; collection 941), realigned to mitigate misalignment around indels on SNV calls (collection 1098), and used to call variants with Naive Variant Caller [24]. Finally, we use Variant Annotator to process VCF datasets generated by Naive Variant Caller and to create a list of variants (collection 1412) and the concatenation tool to reduce collection 1412 into a single table (dataset 1413). This dataset is used for further processing with Jupyter. **C.** The relationship of minor allele frequencies for heteroplasmic sites between tissues (panes A and B) and individuals (panes C and D). **D.** Estimates for bottleneck size with (red) and without (blue) accounting for mitotic segregation.

### 2.4 Summary

The above three examples highlight the power of combining *ad hoc* programmatic analyses with a collection of robust tools already provided by Galaxy. In our opinion this have a potential to revolutionize the ways in which biomedical data analysis is performed. In particular we see the following implications:

1. **Lowering entry barriers.** At this point it is widely acknowledged that every biomedical researcher should be able to at least try performing basic data manipulation and analysis tasks. In practice they are often discouraged from doing this by lack of familiarity with systems such as Jupyter or RStudio and may not know how to configure them for the initial use. Integration of Jupyter into Galaxy gives these users a “risk-free” opportunity to try and learn basic exploratory skills without the need to install or maintain anything.
2. **Allowing re-use and experimentation.** Jupyter notebooks are designed to be shareable, just like Galaxy’s workflows, histories, and datasets. This significantly simplifies re-use: one may, for instance, simply import the notebook we developed in example 3, and apply it to their own data. This also aids in experimentation: what would happen if in the analysis described by [9] we use a different mapper and/or variant caller? It is easy to answer this question by applying the existing notebook to a set of variant calls produced with an alternative workflow.
3. **Increasing collaborative possibilities.** Galaxy is popular with biologists due to the ability to run complex analyses without the need to use the command line interface (CLI). However, this is also the reason why many computational scientists are skeptical and often avoid the platform: they feel constrained without the ability to have the full control of tool execution and workflow construction. Integration of Jupyter will bring the two communities closer: computational scientists and bioinformaticians will be able to develop analyses using interactive environments in the form of notebooks, which will immediately be usable by biomedical researchers.

## 3 Methods

Jupyter integration into Galaxy takes advantage of the recently developed and increasingly popular Docker virtualization platform (https://www.docker.com). It uses the Interactive Environment (IE) plug-in functionality written for Galaxy that also allows integration of other similar tools such as RStudio. The program flow is illustrated in Fig. 3. An IE primarily consists of an Entry Point (IEEP) and an associated configuration file. The IE configuration allows for the setting that all data transfer is done via SSL which is useful for production instances. Additionally, individual sites can specify custom Docker images instead of the default provided Jupyter notebook, allowing administrators to craft docker images more specific to their users. This image will be downloaded and installed from the Docker Hub. The default Docker image is specifically crafted for use in conjunction with the Jupyter Interactive Environment (see below). The IEEP launches a docker container on a random port for communication, and configures it to access Galaxy through environment variables passed to the container. These pieces of information are stored so that Galaxy and the Dockerized Jupyter web service can communicate securely, while isolated from the rest of the Galaxy instance for security reasons. The container is built on top of the official jupyter/minimal-notebook, and provides a Jupyter server, with its dependencies such as NumPy [20], SciPy [21] and Matplotlib [22]. Additionally the image contains several Jupyter kernels (different programming language environments) such as R, Ruby, Haskell, Julia, and Octave. By providing a Docker image with a full suite of scientific analysis tools and libraries, users are able to immediately perform their analysis and calculations. In the Python kernel, additional packages can be installed with the python package manager pip, the same is true for the other kernels with their associated package managers. Moreover, tools that can be installed and run in a non-privileged user account can be added to the container on demand. Once the container has launched on the backend, it is visually embedded inside the Galaxy interface, at which point it can be used to interactively program, develop, and analyse data in any of the aforementioned programming languages. During Docker container startup, a cron job is launched, monitoring whether the IE is still being used by checking the network traffic. If the page with the Interactive Environment is closed, the Docker process automatically terminates itself. Within the Jupyter Notebook, two important custom functions are defined which enable the user to load data from the history or store data to the Galaxy history using the Galaxy API [23]. The get function expects one parameter, the identifier of the dataset as shown in the history. The put function automatically builds a connection to the host Galaxy instance and transfers a specified file from inside the Docker container to the user’s history. The flexibility is obvious: any dataset can be loaded into the notebook, datasets can be combined or modified, and the results can be written back to the history. The entirety of Galaxy-IE communication occurs between the Galaxy host and the Docker container, without the need for the user to upload or download data to their personal workstation. This is not only faster in most cases it has also positive implications on data security as the data did not leave the compute center.

**Figure 3.**
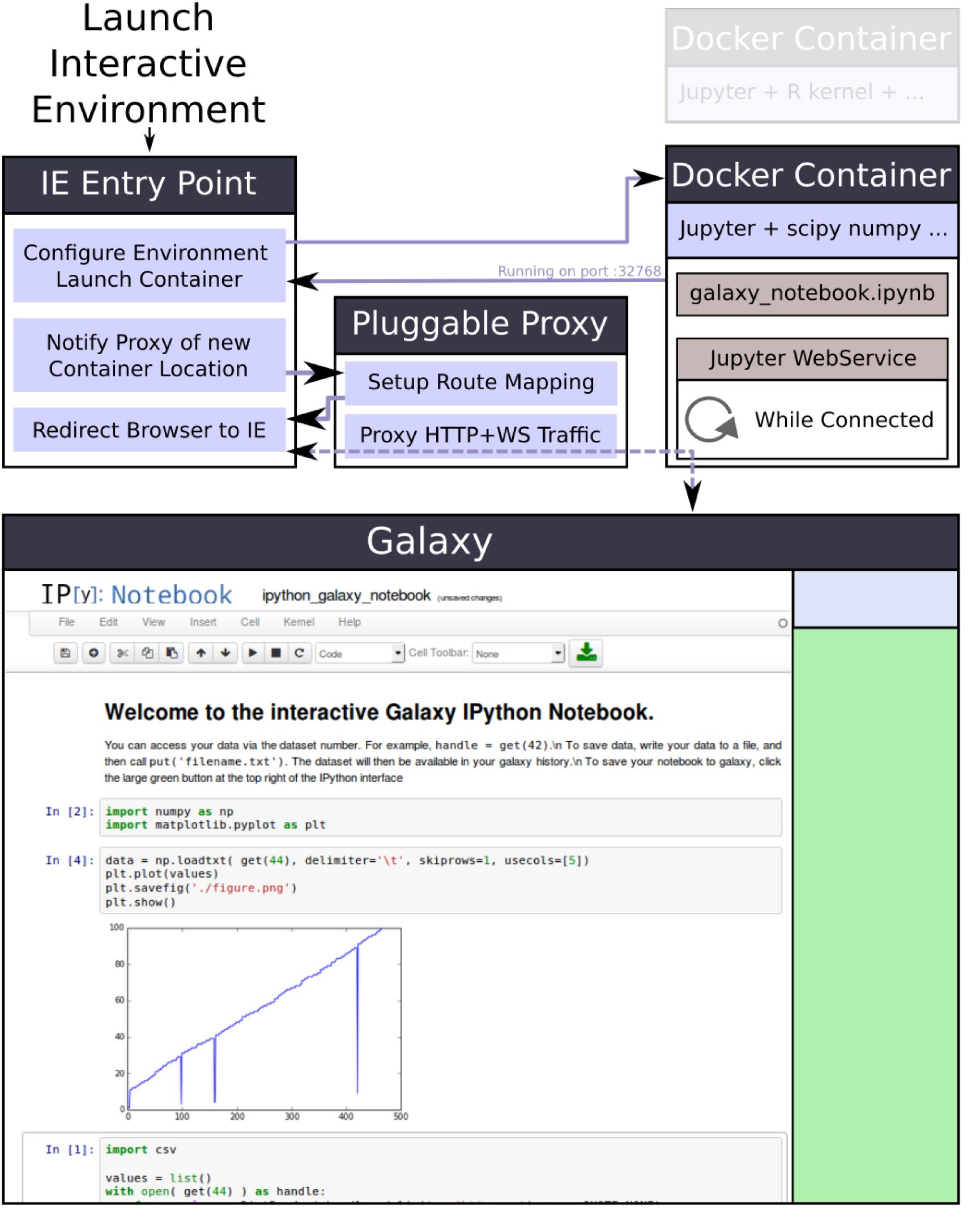
Schematic representation of Galaxy/Jupyter architecture taking advantage of Docker functionality.

Notebooks can be saved to the Galaxy history at any time; once in Galaxy’s history they can be inspected like any other Galaxy dataset, allowing for a read-only view of the analysis steps which we run. Additionally, notebooks can be reused. A new Jupyter instance is created that retains the stored work. This functionality ensures the reproducibility of data analysis and is therefore an essential feature of the Jupyter Interactive Environment.

## 4 Acknowledgements

We are grateful to the members of Galaxy development team for their help with preparation of this manuscript. All experimental procedures used to generate datasets used in Example 3 comply with the principles of the Helsinki Declaration. All data has been collected under IRB Protocol No.30432EP. In this study patients at the pediatric outpatient clinic (located the Pennsylvania State University Hershey Medical Center in Harrisburg, Pennsylvania) were approached by experienced coordinators and invited to participate. Samples were collected from participants providing verbal consent in the clinic. Only date of birth and date of collection were associated with each sample. No personally identifiable information was recorded. This project was supported by NIH/NHGRI Grant U41 HG005542 (JT and AN), a German Federal Ministry of Education and Research grant 031A538A de.NBI (RB), and Collaborative Research Centre 992 Medical Epigenetics grant SFB 992/1 2012 (RB). Additional funding was provided by Huck Institutes for the Life Sciences at Penn State and, in part, under a grant with the Pennsylvania Department of Health using Tobacco Settlement Funds. The Department specifically disclaims responsibility for any analyses, interpretations or conclusions.

## 5 Supplemental materials

**Supplemental Figure 1**. Schematic representation of Galaxy/Jupyter architecture taking advantage of Docker functionality.

**Supplemental Document 1**. A PDF version of Jupyter notebook for Example 1: Visualization of read coverage across HIV genome.

**Supplemental Document 2**. A PDF version of Jupyter notebook for Example 2: Read count normalization and variance shrinkage for an RNA-seq experiment.

**Supplemental Document 3**. A PDF version of Jupyter notebook for Example 3: Analysis of heteroplasmy dynamics in humans.

